# MAW - The Reproducible Metabolome Annotation Workflow for Untargeted Tandem Mass Spectrometry

**DOI:** 10.1101/2022.10.17.512224

**Authors:** Mahnoor Zulfiqar, Luiz Gadelha, Christoph Steinbeck, Maria Sorokina, Kristian Peters

## Abstract

Mapping the chemical space of compounds to chemical structures remains a challenge in metabolomics. Despite the advancements in untargeted liquid chromatography-mass spectrometry (LC-MS) to achieve a high-throughput profile of metabolites from complex biological resources, only a small fraction of these metabolites can be annotated with confidence. Many novel computational methods and tools have been developed to enable chemical structure annotation to known and unknown compounds such as *in silico* generated spectra and molecular networking. Here, we present an automated and reproducible <u>M</u>etabolome <u>A</u>nnotation <u>W</u>orkflow (MAW) for untargeted metabolomics data to further facilitate and automate the complex annotation by combining tandem mass spectrometry (MS^2^) input data pre-processing, spectral and compound database matching with computational classification, and *in silico* annotation. MAW takes the LC-MS^2^ spectra as input and generates a list of putative candidates from spectral and compound databases. The databases are integrated via the R package Spectra and the metabolite annotation tool SIRIUS as part of the R segment of the workflow (MAW-R). The final candidate selection is performed using the cheminformatics tool RDKit in the Python segment (MAW-Py). Furthermore, each feature is assigned a chemical structure and can be imported to a chemical structure similarity network. MAW is following the FAIR (Findable, Accessible, Interoperable, Reusable) principles and has been made available as the docker images, maw-r and maw-py. The source code and documentation are available on GitHub (https://github.com/zmahnoor14/MAW). The performance of MAW is evaluated on two case studies. MAW can improve candidate ranking by integrating spectral databases with annotation tools like SIRIUS which contributes to an efficient candidate selection procedure. The results from MAW are also reproducible and traceable, compliant with the FAIR guidelines. Taken together, MAW could greatly facilitate automated metabolite characterization in diverse fields such as clinical metabolomics and natural product discovery.

## 1 Introduction

Over the last few decades, omics technologies have gained importance in the biological domain. One of the recent developments in omics science is metabolomics, which is a collection of high-throughput analytical methods to assess the metabolic profiles of biological samples [1]. Among metabolomics approaches, liquid chromatography coupled to mass spectrometry (LC-MS) is the most often-used technique for the identification of lower molecular weight (typically below 1000 Da) molecules. LC-MS also provides broad coverage of biologically relevant metabolites [2] or natural products [3]. However, as metabolomics data can be complex and convoluted, chemical heterogeneity poses a challenge to the number of metabolites that can be identified with metabolomics [4]. Untargeted metabolomics data, which represents global metabolic features in a sample, results in a large number of files containing spectral information from thousands of metabolite features [5]. Unlike genomics where there are only 4 nucleotides and finite possible combinations, chemical structures are made up of many constituents leading to an almost infinite chemical space. In addition, it is also difficult to reproduce the same metabolic features between different instruments, which makes the structural identification of the metabolic features difficult [6].

Typically, an LC-MS metabolomics pipeline involves the following steps: (1) conversion of RAW spectral data to an open-source machine-readable format such as .mzML, (2) peak detection, (3) peak alignment, (4) retention time correction, (5) isotope detection, (6) adduct annotation, (7) peak annotation, and (8) data analysis [7–9]. To acquire more structural information about the samples, tandem mass spectrometry is performed, where two (MS^2^) or more (MS^n^) analyzers are combined with a collision chamber which performs fragmentation of the peaks picked in MS^1^ [10]. The fragmentation spectra from MS^2^ are then used to perform metabolite annotation. To provide a distinction between different levels of confidence for these annotations, the Metabolomics Standards Initiative (MSI), in 2007, proposed four levels of identifications [11] which were later expanded [12] to five levels. The terms annotation and identification are used interchangeably. However, the term annotation refers to the putative association of a compound to a structure, as compared to the term identification, where the identity of a compound is confirmed with a given metabolic standard. These levels of identifications are described in Table 1 in the results section.

**Table 1:**
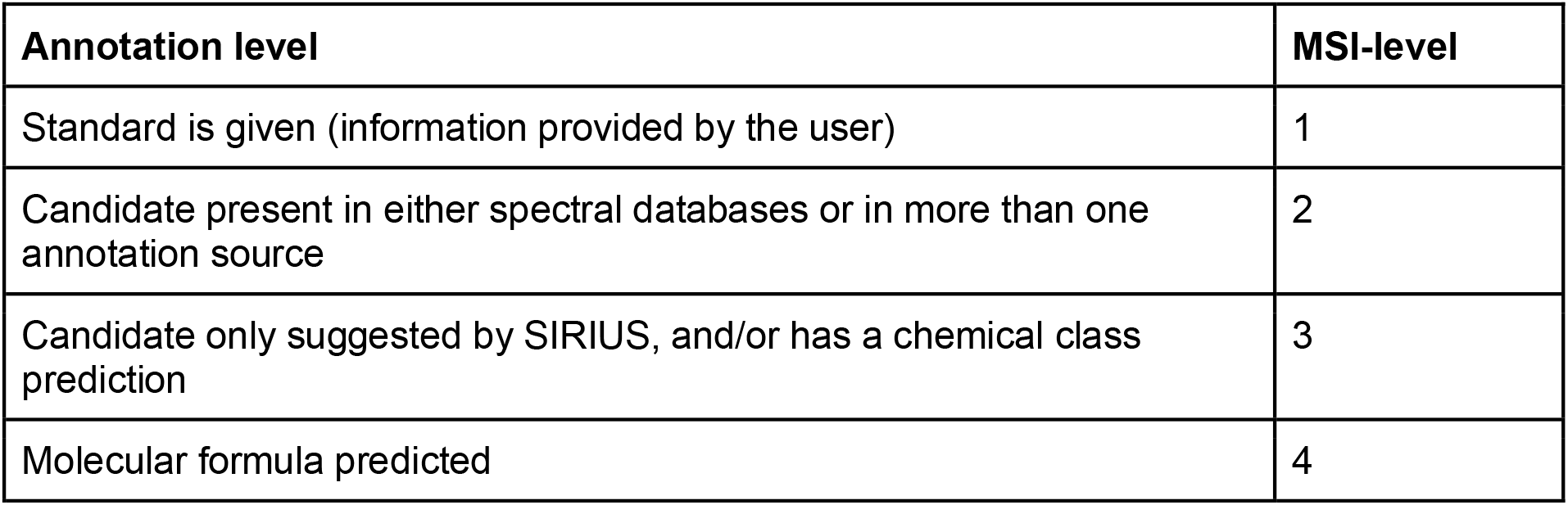
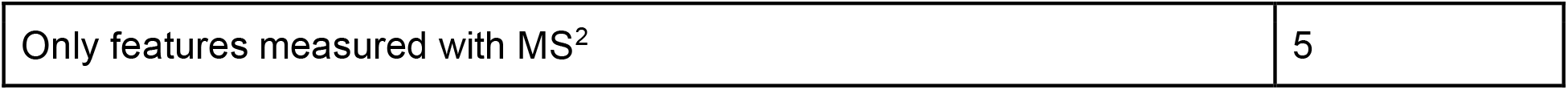
MAW operates on MSI-levels [12] and categorizes annotations based on the rules given above.

To support metabolomics identification, different data sources such as spectral, compound, and pathway databases have been integrated into metabolomics data processing pipelines and routines [13]. To this end, many software tools and programmatic approaches such as R packages have been developed for metabolite annotation that use LC-MS^2^ as input data. A few of the most popular examples are SIRIUS [14], and MetFrag [15]. Some other relatively new tools are MetaboAnnotatoR [16],, CFM-ID 4.0 [17], DEREPLICATOR(+) [18], molDiscovery [19], CluMSID [20], and MS2Query [21]. These tools are termed annotation tools. They generally annotate compounds based on spectral and compound libraries and databases. However, many tools currently only partly support the FAIR [22] criteria, specifically the reusibility guidelines. Some of these tools have already been integrated into computational metabolomics workflows such as PhenoMeNal [23], Workflow4Metabolomics (W4M) [24], or GNPS (Global Natural Products Social Molecular Networking) [25]. These tools utilize workflow management systems, such as Galaxy [26], KNIME [27], TidyMass [28], or work as independent software such as MS-Dial [29], or mzMine [30] which can be integrated into any metabolomics pipeline or workflow..

Customization is a very important aspect of these metabolomics workflows due to the variations in the experimental designs and samples. Annotation, for instance, is also usually a manual process to select the best candidate by going through the candidate list provided by different annotation tools, which takes effort and time, specifically in the case of untargeted metabolomics, which is a comprehensive analysis of all measured compounds in a sample. Untargeted metabolomics studies, and phenome centers, where large amounts of data are generated repeatedly, result in large-scale data where workflows can greatly automate and facilitate the analysis [31, 32]. For such large datasets, reproducibility [33] can be another issue where the intermediate results should be tracked to check the final outcome. In other words, the provenance [34] should be recorded. In addition to reproducibility, best practices defined in the FAIR principles [22, 35] should be employed within workflow systems to improve the Findability, Accessibility, Interoperability, and Reusability of the research objects.

Here, we introduce the reproducible and automated <u>M</u>etabolome <u>A</u>nnotation <u>W</u>orkflow (MAW) for untargeted metabolomics data. MAW is a computational workflow which can automate execution in e.g., cloud environments and can greatly speed up the analysis of high-throughput data. MAW performs (1) MS^2^ data preprocessing, (2) spectral database dereplication, (3) compound database dereplication, (4) candidate selection, and (5) chemical compound classification using various tools integrated into the workflow. It provides a list of top-scoring candidates as a network and also prioritizes a final candidate from the list, for each feature. MAW follows the FAIR guiding principles [35]. It uses public databases and open-source software tools. The complete MAW source code is available on GitHub [36]. It is developed in the languages R and Python and can be executed with all its functionalities and dependencies as two docker images for MAW-R and MAW-Py from DockerHub [37, 38].

## 2. Methods

### 2.1 Input Data Prerequisites

The objective of this workflow is metabolome annotation from the data acquired from liquid chromatography-tandem mass spectrometry (LC-MS^2^) with data-dependent acquisition (DDA) mode [39]. MAW takes .mzML files as input \containing both MS1 features and MS^2^ spectra. The LC-MS features should be pre-selected, as MAW directly starts with the pre-processing of MS2 spectra for annotation. The RAW files after data acquisition can be converted to .mzML format [7] using the MS-Convertor from the ProteoWizard suite [40] or the mass spectrometry file conversion tool on GNPS. The profile peak data should be converted to centroid peak data during the conversion.

We used two reference datasets to assess the performance of MAW. The first dataset consists of standards from a study on hypersalinity in diatoms [41]: betaine, pipecolinic acid, cysteinolic acid, methionine sulfoxide, N,N-dimethylarginine, O-acetyl-L-carnitine, O-propanoyl-L-carnitine, O-butanoyl-L-carnitine, and isovalerylcarnitine. The second dataset is from a study on nine different Bryophyte species [42]. The .mzML files of the entire study on Bryophytes are available in MetaboLights under the accession MTBLS709. Both datasets were obtained using LC-MS^2^ in DDA mode. The mzML files used in this study have been made available on Zenodo [43, 44].

To use MS^2^ Spectral databases in MAW, we downloaded GNPS, HMDB (Human Metabolome Database) [45], and MassBank [46]. These databases were stored as an R object using MsBackend, which is a virtual class defined in the R package Spectra [47] to store and retrieve the mass spectrometry data. The database dumps have been made available on Zenodo [48].

### 2.2 Metabolome Annotation Workflow

The Metabolome Annotation Workflow (MAW) is executed as a computational workflow which provides different executable modules: 1) Spectral database dereplication, 2) Compound database dereplication, and 3) Candidate selection. MAW is written in R for metabolomics data handling and Python for cheminformatics analysis. Figure 1 illustrates the overview of the workflow.

**Figure 1:**
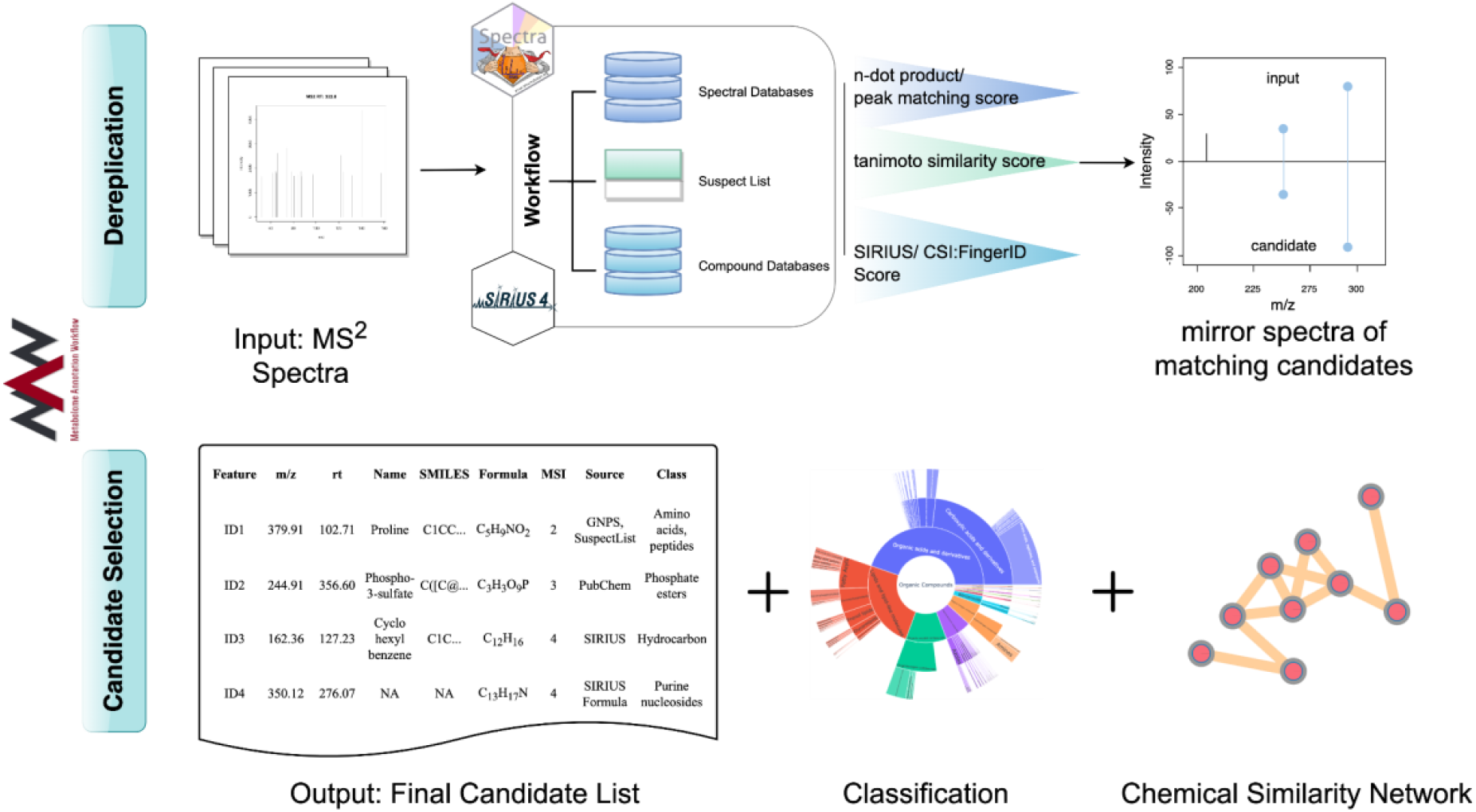
Overview of MAW. MAW-R executes dereplication. It starts with the input data files and searches the spectra in spectral and compound databases with the R package Spectra and SIRIUS respectively. Once the dereplication is completed and each file and precursor mass [m/z] has some candidates from different databases, MAW-Py performs candidate selection from these sources, generates classes, and creates a table for the chemical similarity network, which can be visualized in Cytoscape [49].

#### 2.2.1 MAW-R Segment

The workflow starts with preprocessing MS^2^ spectra using the package Spectra. The function spec_Processing in MAW-R, reads the .mzML files and extracts all precursor masses [m/z]. This step is common for both spectral and compound database dereplication. Figure 2 describes the MAW-R segment in a stepwise manner.

**Figure 2:**
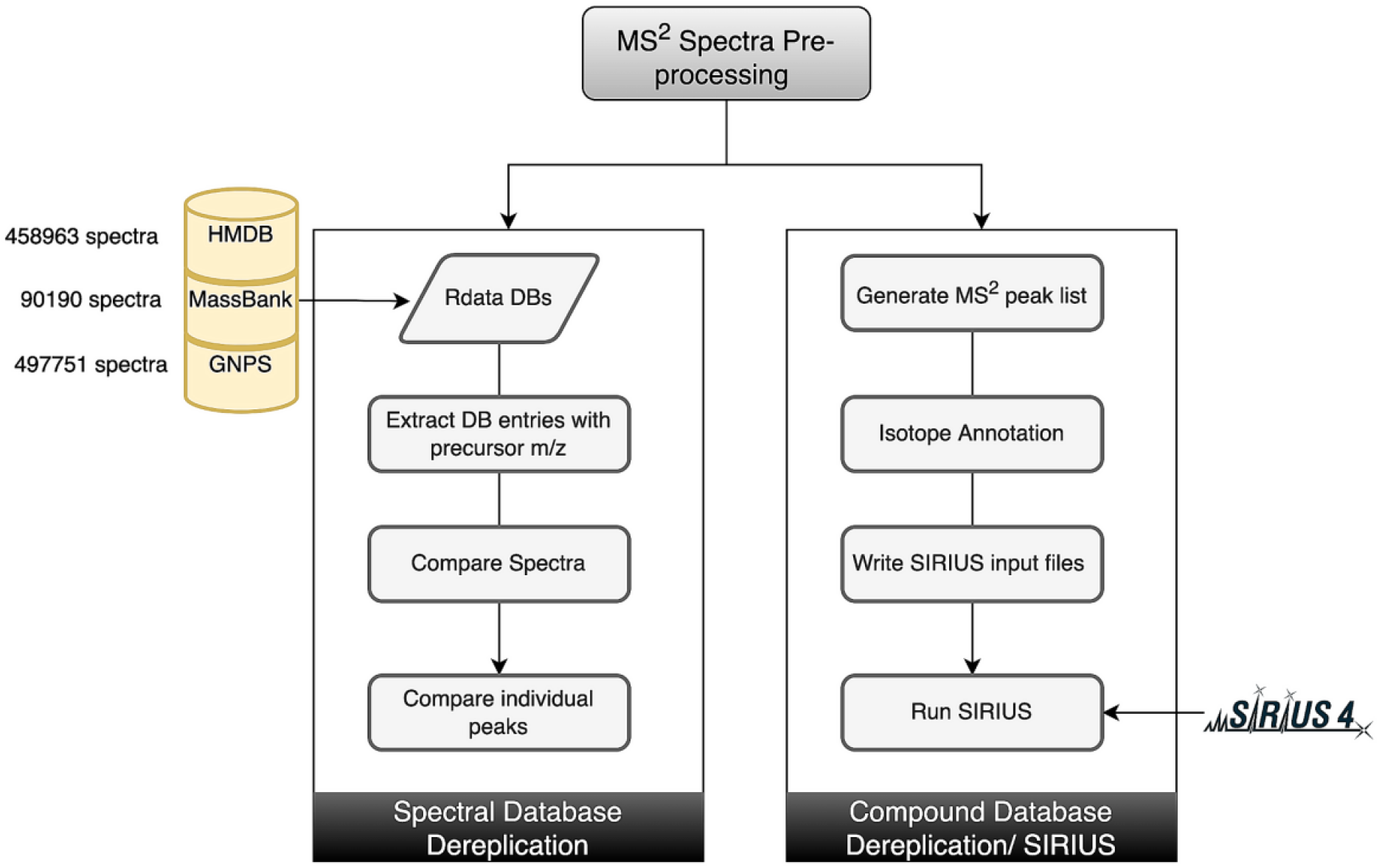
Overview of spectral and compound database dereplication performed in MAW-R. Spectral database dereplication utilizes HMDB, MassBank, and GNPS for each precursor mass [m/z]. The spectral databases are stored as Rdata objects, available on Zenodo. The spectra from input queries and databases with precursor mass [m/z] of interest are extracted. These spectra are compared with cosine similarity scores between the query and database spectra. Individual peaks are also compared to remove any false positive candidates with high overall similarity but lower individual peak similarity. The second part of MAW-R is the compound database dereplication with SIRIUS CLI. To prepare the input files for SIRIUS, MS^2^ peaks and isotopic peaks are extracted. This information is written into SIRIUS input files (.ms), which are used to run SIRIUS.

##### 2.2.1.1 Spectral Database Dereplication

To start spectral database dereplication, the `spec_dereplication_file` function is used. This module requires GNPS, HMDB (in silico and experimental spectra), and MassBank spectra stored as R objects. The first function within this module is the `spec2_Processing` function which extracts all spectra with a given precursor mass [m/z] value, normalizes the intensity, and removes low-intensity peaks (<5%). This function is applied to both query and database candidate spectra so that they are comparable.

GNPS uses different scoring as compared to HMDB and MassBank. In order to integrate both types of scoring, we implemented the following approach. For GNPS, the fragment peaks of query and candidate spectra are defined as matching peaks, if the difference between their fragment mass [m/z] values and the difference between these adjusted fragment mass [m/z] values is less than the given tolerance (15 ppm by default). This scoring parameter is set to 0.85 (maximum = 1.0) as the threshold. For HMDB and MassBank, a comparison of the two spectra with the dot product is used to determine matching peaks. The default is set to 0.70.

The top candidates are also given a normalized score of similarity between fragment masses [m/z] of individual matching peaks and the intensity of these peaks by function `peakdf`. The ratio of matching peaks by the total peaks in query spectra is also weighed within these scores.

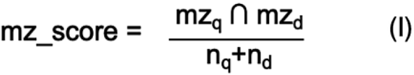

The mz_score is the intersection of the matching peaks between two comparing spectra. mz_q_ ∩ mz_d_ expression defines the number of matching peaks. n_q_ is the total number of peaks in the query spectrum and n_d_ is the total number of matching peaks in the database spectrum.

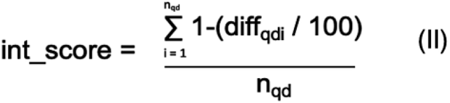

The int_score is the average summation of the difference between individual matching peaks defined by diff_qdi_. Since the score is normalized to 1, the diff_qdi_ values are divided by 100 and then subtracted from 1 to give more weightage to peaks with a lesser difference in intensity.

##### 2.2.1.2 Compound Database Dereplication

To utilize the Command Line Interface (CLI) of SIRIUS (version 4.9.12), a parameter file is required as an input for each precursor mass [m/z]. The `sirius_param` function is used to generate these parameter files in SIRIUS readable .ms format. The parameter file contains the id of the feature, precursor mass [m/z], charge, the median of retention time [sec], isotopic peaks if present, and the MS^1^ and MS^2^ peaks. In order to extract this information, `ms2_peaks` and `ms1_peaks` functions are required. `ms2_peaks` extracts and combines the fragment peak lists for each precursor mass [m/z]. The R package CAMERA is used to extract MS^1^ peaks and perform isotope annotation. To run SIRIUS, the `run_sirius` function is applied based on the isotopic peak annotation. Here, the user can define the database to be used within SIRIUS. For the standards dataset, the “all” database was used, which is a combination of many databases such as PubChem, HMDB, and COCONUT (COlleCtion of Open Natural ProdUcTs). The “all” database is also the default setting. For the bryophytes dataset, the “bio” database was used, which is a collection of different chemical databases reserved for biological sources. Another database used for annotating the bryophytes dataset is the COCONUT database [50]. For compound classification, CANOPUS is used [51]. Figure 2 describes the step-wise functions in the compound database dereplication module, using SIRIUS.

#### 2.2.2 MAW-Py Segment

##### 2.2.2.1 Post-processing

To resolve missing SMILES (Simplified Molecular Input Line Entry Specification) and non-conventional compound names from GNPS, the function `spec_postproc` removes any naming anomalies and adds SMILES using PubChemPy [52]. For HMDB, the results only contain the HMDB database ID. To obtain the names, formulae, and SMILES, the structures.sdf file is downloaded from HMDB. Using the same function, any spectral match with mz_score, int_score, and the ratio of matching peaks below 0.50 are discarded. To post-process the results from SIRIUS, the function `sirius_postproc` adds the top-scoring candidates from SIRIUS. If there are no structure candidates predicted, then the function adds the top-ranking formulae as the candidates.

##### 2.2.2.2 Candidate Selection

The candidate selection is performed by the function `CandidateSelection_SimilarityandIdentity`, which uses RDKit [53] to perform the calculations. takes results from spectral and compound databases and runs the Tanimoto similarity score (keeping a threshold of >= 0.85) to search for common structures or substructures among the top predicted candidates from all these sources. The results can be visualized in chemical similarity networks. This provides the most probable candidate structure. To select one candidate among these top candidates, without manually visualizing the network or going through the list of top candidates, further calculations are performed, with a Tanimoto similarity score of >= 0.99. This calculation selects one candidate that is present among the top candidates from all or most of the sources. In case of a disagreement among the sources, we provide a prioritization scheme with results from GNPS given higher priority, as compared to SIRIUS, MassBank, and HMDB (see result section 3.2). The result is represented by a list of features and corresponding candidates with MSI confidence levels of identification. Figure 3 explains the step-wise processing of the candidate selection. To visualize the 2D structures, CDK-Depict [54, 55] can be used, as was used in the depiction of structures in Table 2, Table 3 and Table 5.

**Figure 3:**
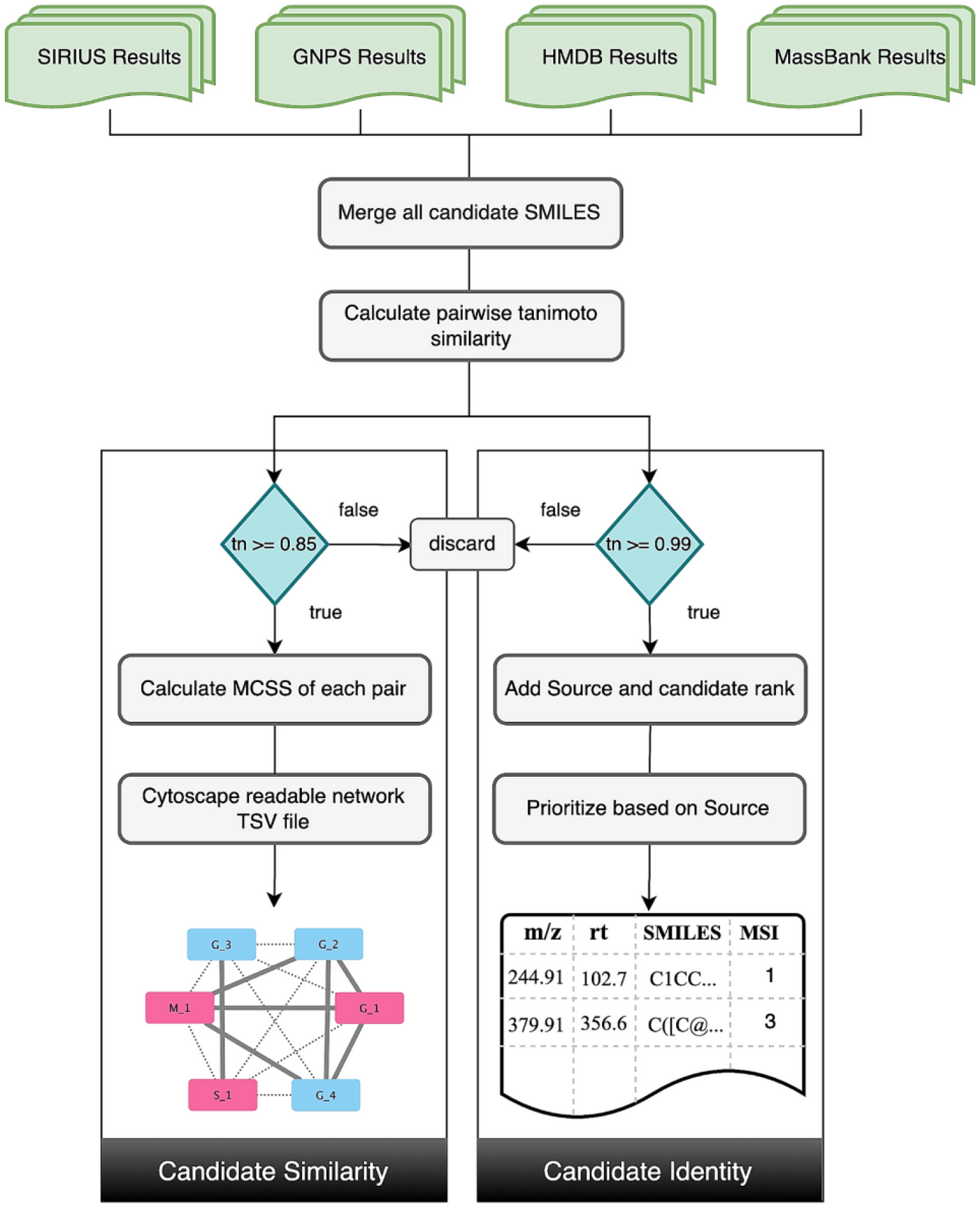
Overview of the candidate selection performed within MAW-Py. The workflow takes the results of top-scoring candidates from all sources and merges the list of top-scoring SMILES from these sources in a single CSV file. These SMILES are then searched for structural similarity. For the pairwise “Chemical Similarity” calculation part, the SMILES of candidate compounds with Tanimoto similarity scores of < 0.85 when compared with other SMILES in the list of top scoring candidates, are discarded. For the rest of the SMILES, a Maximum Common Substructure (MCSS) is calculated for each pair of SMILES in the list. Based on this MCSS calculation, a TSV file is generated which can be used in Cytoscape to visualize the chemical similarity among the candidates. Candidates that belong to a cluster with the most number of candidates from top ranks should be considered as the most probable structures and substructures for the particular feature. For the “Candidate Identity” part, the threshold is >= 0.99. The candidate identity among SMILES leads to a single structure as the top-ranking candidate for each feature. If there is no identity, or the sources provide different top-ranking structures, a prioritization is performed. In total, there are four sources (SIRIUS, GNPS, HMDB, and MassBank). The scheme is – three sources with the same candidate > two sources with the same candidate > single source (GNPS) > single source (SIRIUS) > single source (MassBank or HMDB). The scheme is defined from the results obtained with known compounds from the standards dataset from diatoms.

**Table 2:**
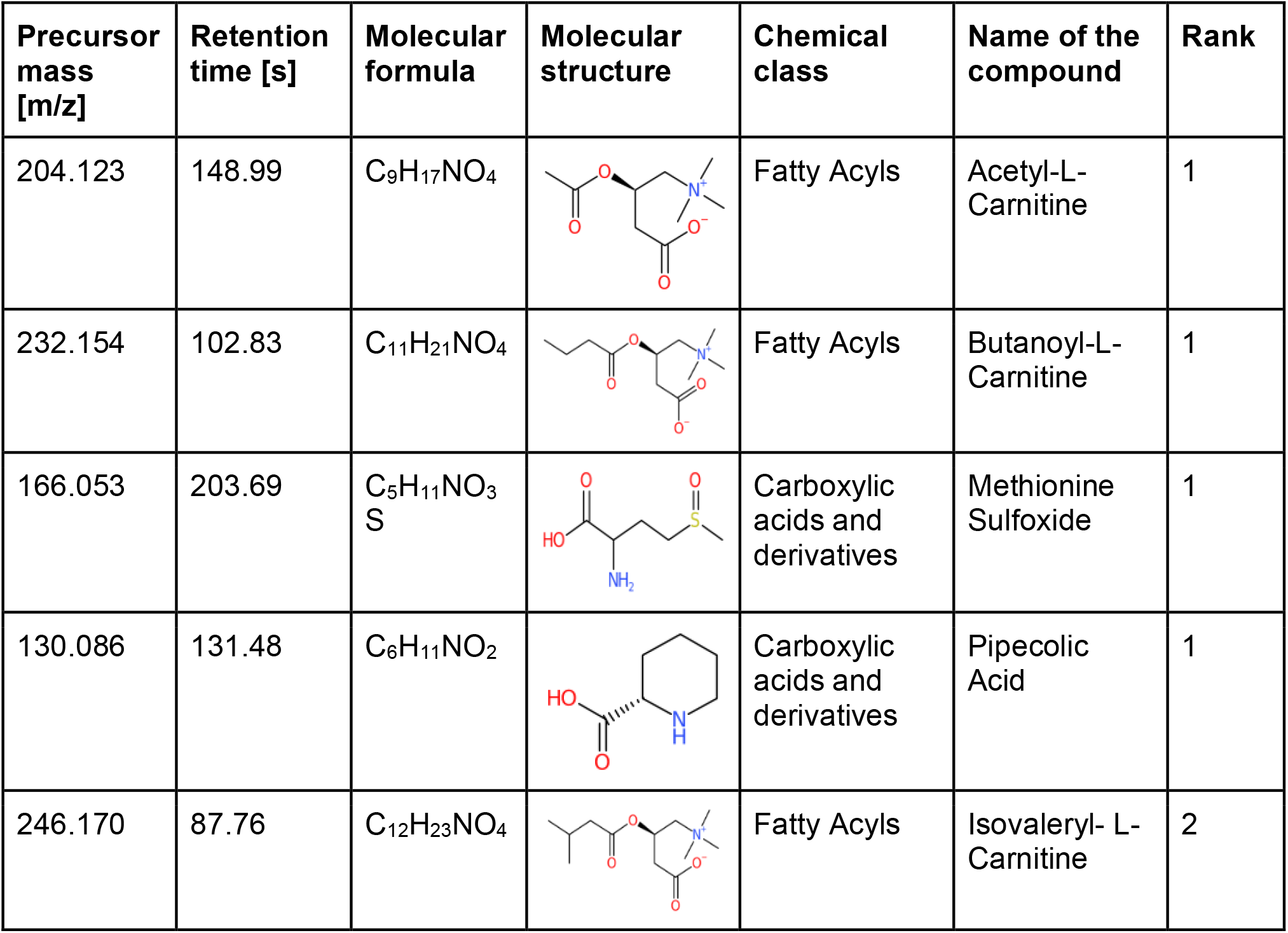
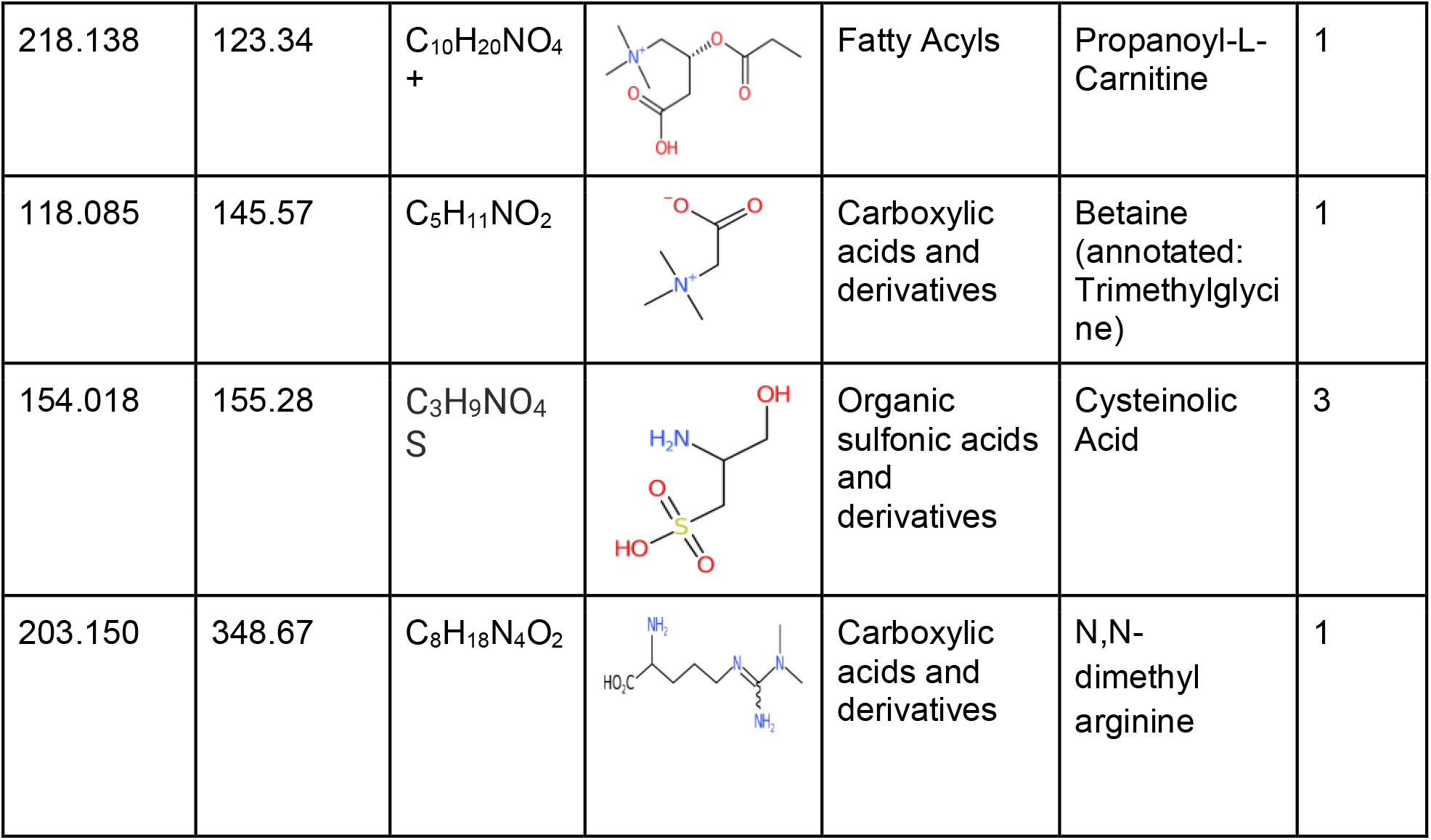
List of molecules from the standards dataset and top candidates from MAW. The precursor mass [m/z] and retention time [s] represent the mass of the precursor ions given in mass-to-charge ratio and the median retention time of the precursor ion given in seconds, respectively. The given ranks are from MAW.

**Table 3:**
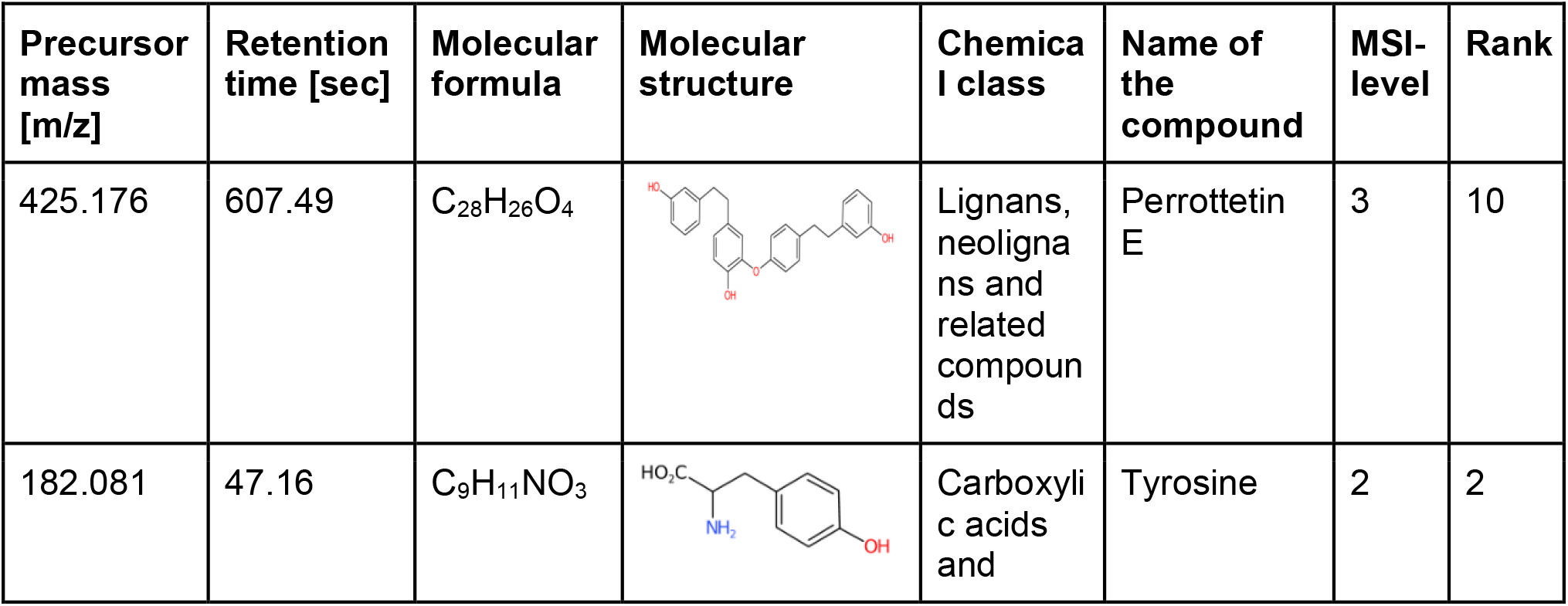
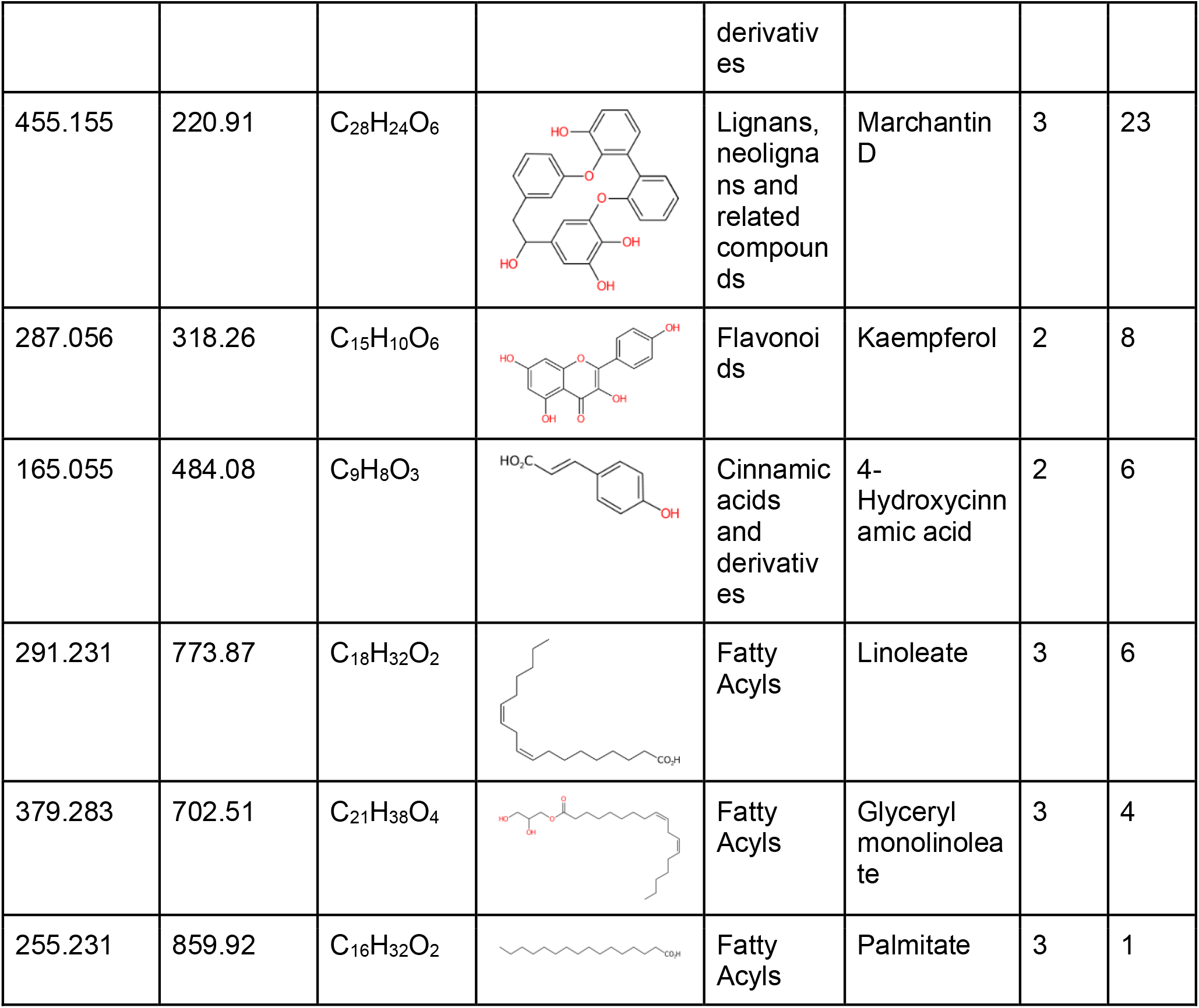
Features that were confirmed using MAW and the original study on the bryophytes data. The precursor mass [m/z] and retention time [s] represent the mass of the precursor ions given in mass-to-charge ratio and the median retention time of the precursor ion given in seconds, respectively. The molecular formula and name of the compound are extracted using PubChemPy or SIRIUS, depending on the annotation source. The chemical classes of the compounds are either extracted from ClassyFire or CANOPUS. All the molecular structures are generated via CDK-Depict. The MSI-levels are in accordance with the rules described in Table 1. The given ranks are from MAW, using the database COCONUT in the compound dereplication module.

| Precursor mass [m/z] | Retention time [sec] | Molecular formula | Molecular structure | Chemical class | Name of the compound | MSI-level | Rank |
| --- | --- | --- | --- | --- | --- | --- | --- |
| 425.176 | 607.49 | $C_{28}H_{26}O_4$ | | Lignans, neolignans and related compounds | Perrottetin E | 3 | 10 |
| 182.081 | 47.16 | $C_9H_{11}NO_3$ | | Carboxylic acids and derivatives | Tyrosine | 2 | 2 |
| 455.155 | 220.91 | C<sub>28</sub>H<sub>24</sub>O<sub>6</sub> |  | Lignans, neolignans and related compounds | Marchantin D | 3 | 23 |
| 287.056 | 318.26 | C<sub>15</sub>H<sub>10</sub>O<sub>6</sub> |  | Flavonoids | Kaempferol | 2 | 8 |
| 165.055 | 484.08 | C<sub>9</sub>H<sub>8</sub>O<sub>3</sub> |  | Cinnamic acids and derivatives | 4-Hydroxycinnamic acid | 2 | 6 |
| 291.231 | 773.87 | C<sub>18</sub>H<sub>32</sub>O<sub>2</sub> |  | Fatty Acyls | Linoleate | 3 | 6 |
| 379.283 | 702.51 | C<sub>21</sub>H<sub>38</sub>O<sub>4</sub> |  | Fatty Acyls | Glycerol monolinoleate | 3 | 4 |
| 255.231 | 859.92 | C<sub>16</sub>H<sub>32</sub>O<sub>2</sub> |  | Fatty Acyls | Palmitate | 3 | 1 |

##### 2.2.2.3 Compound Classification

Annotation for compound classes can be achieved by a function called `classification`. CANOPUS is a chemical classification prediction tool, integrated in SIRIUS and annotates a chemical class using the LC-MS^2^ spectra. To provide classification to annotations that have an annotation source other than SIRIUS (such as GNPS, HMDB, or MassBank), ClassyFire [56] is also integrated into the workflow which takes SMILES as input. Both CANOPUS and ClassyFire are based on the ChemOnt ontology [56]. To use ClassyFire in Python, the pybatchclassyfire [57] package is used.

### 2.3 Towards a FAIR and Reproducible MAW

Computational workflows like MAW can be described as digital research objects in their own right [58]. As a result, the FAIR (Findable, Accessible, Interoperable, and Reusable) principles can be applied to these workflows. For good scientific practice, in MAW, we implemented the recommended FAIR principles. To enable Findability and Accessibility, the source code of MAW has been made available in the GitHub repository zmahnoor14/MAW. In addition, MAW has been versioned through Zenodo-DOIs generated from the GitHub repository. To enable Interoperability and Reusability, separate Docker images are available on DockerHub for MAW-R and MAW-Py. In *Future Developments* in the discussion subsection, we detail additional practices that will be implemented to further support reproducibility and the FAIR principles.

## 3 Results

### 3.1 Annotations from Tandem Mass Spectrometry Data

To evaluate the performance of MAW, we utilized two datasets, a list of standard molecules from the diatoms (further called the standards dataset), and untargeted LC-MS^2^ spectra from the bryophytes dataset (further called the bryophytes dataset). Each high-scoring candidate is given MSI-level of confidence [12]. A detailed overview of these MSI-levels is presented in Table 1.

The MSI-levels 1 and 2 represent known compounds that are present in spectral databases (see Table 1). In the case of MSI-level 1, an analytical standard is utilized to confirm their identification. Hence, results from the standards dataset are used to validate the results from MAW. Out of 9 standard metabolites, 6 were correctly identified as the top first candidates with MAW as shown in Table 2. Three particular cases with second or third ranks are: (a) the reduced form of betaine was annotated as the top candidate (trimethylglycine). (b) Cysteinolic acid wasn’t detected in spectral databases, rather 1-chlorobenzotriazole was annotated by both GNPS and MassBank. Only SIRIUS identified cysteinolic acid as the top candidate. According to our current annotation source prioritization scheme (described in Figure 3), 1-chlorobenzotriazole from GNPS and MassBank is at the top two ranks, and cysteinolic acid from SIRIUS received an overall rank of 3 in MAW. (c) Valeryl L-carnitine was detected as the top candidate by GNPS and MassBank. In contrast, the correct compound isoveleryl-L-carnitine was detected as the top candidate from SIRIUS and the second top candidate from GNPS. Some correct matches might have been discarded due to the scoring threshold. For example, butanoyl carnitine search in MassBank had no results because the threshold set for MassBank score is 0.70. The correct annotation for this compound within MassBank had a score of 0.44, which is lower than the set threshold.

In the bryophyte dataset, 48 features were originally annotated with chemical structures, and 604 features were originally assigned with chemical classes using the MetFamily classifier [59]. Using MAW, when choosing the “bio” and COCONUT databases for structural annotation in SIRIUS, we confirmed a total of 8 compounds similar to the original study (see Table 3). Using the COCONUT database in SIRIUS, gave slightly better ranks than the “bio” database. In total, using MAW, 881 out of 933 features were annotated with chemical structures when compared to 48 structurally annotated features of the original study. For 92 features, MAW provided only a compound class and a formula, and for 18 features only a formula. These results are observed with default settings in MAW.

To classify MS^2^ spectra into compound classes, MAW used CANOPUS and ClassyFire (ChemONT ver 2.1). We found the most prevalent superclasses to be Lipids and lipid-like molecules, Organic acids and derivatives, Benzenoids, and Phenylpropanoids and polyketides. The original study found the superclasses Lipids and lipid-like molecules, Phenylpropanoids and polyketides, Lignans, neolignans and related compounds, and Organic oxygen compounds to be the most diverse but the authors used a different classifier. Figure 4 shows a sunburst plot providing an overview of the annotated superclasses, classes, and subclasses identified in MAW.

**Figure 4:**
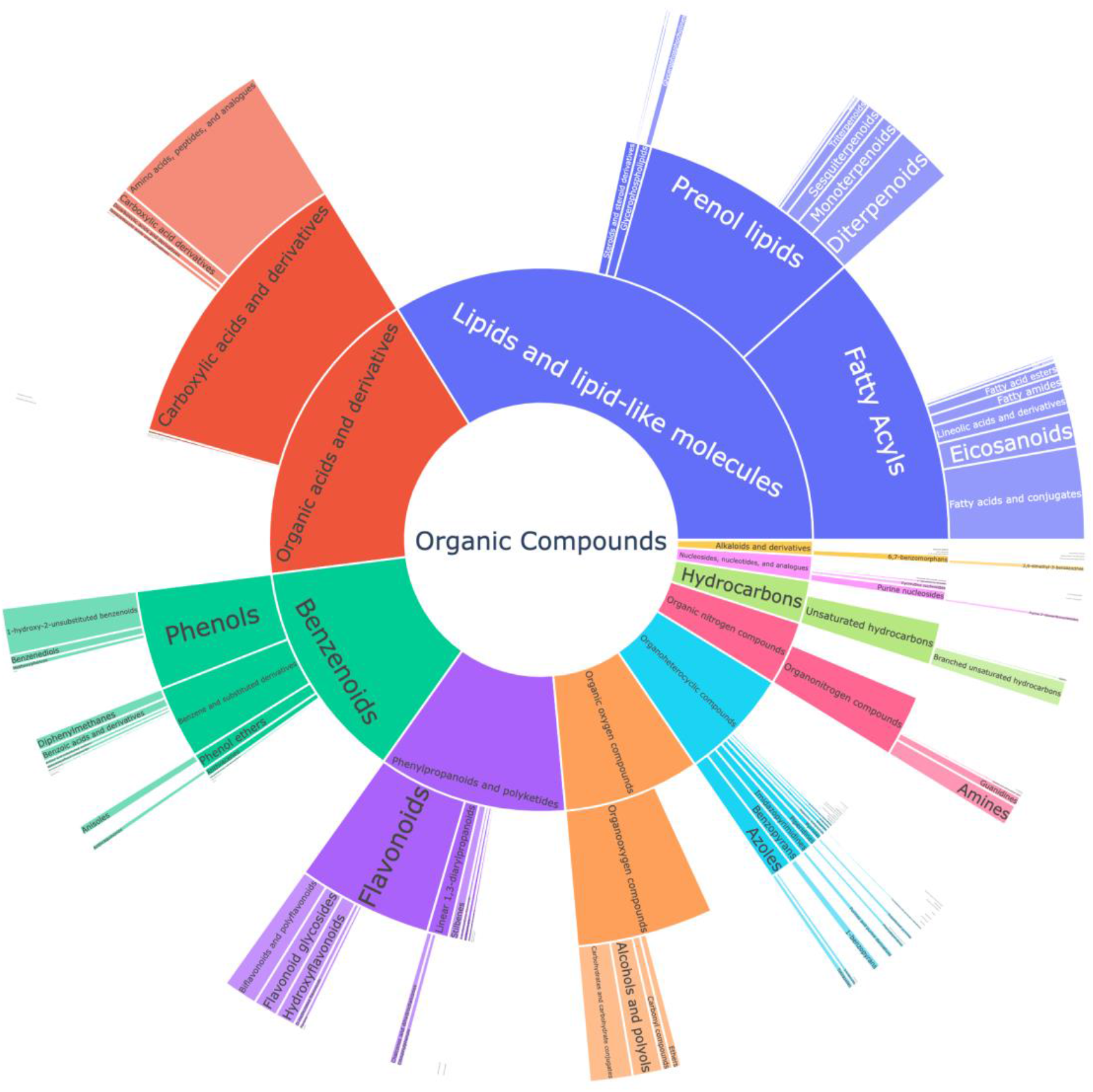
The sunburst plot showing the chemical diversity of the classified top-ranking annotations. Each shown compound class is represented by the counts of classified compounds belonging to the respective class. In the center, the most general superclasses are shown. The further to the outside, the more specific compound classes are shown. MAW uses CANOPUS and ClassyFire to annotate chemical classes to features. MAW found the subclasses Lipids (33.92%), Organic acids (17.02%), Benzenoids (13.94%), and Phenylpropanoid and polyketides (11.34%) to be most abundant.

### 3.2 Comparison of MAW with integrated annotation tools using the standards dataset

We evaluated MAW’s performance with regard to the integrated annotation tools SIRIUS, and Spectra (GNPS, Massbank, and HMDB) using the standards dataset. MAW identified 8 out of 9 compounds correctly except for Betaine (where the annotated compound is the reduced form of Betaine). Isovaleryl carnitine ranked second and cysteinolic acid ranked third, as shown in Table 2. In the case of using SIRIUS without spectral databases, fatty acyls (except for Butanoyl Carnitine) were annotated as isomers or similar structures instead of the actual compounds. SIRIUS correctly annotated methionine sulphoxide, N,N-dimethylarginine, and carboxylic acid and its derivatives. GNPS correctly annotated all compounds except for betaine and cysteinolic acid. MassBank and HMDB showed the lowest performances. MassBank only annotated N,N-dimethyl arginine, and acetylcarnitine with a score threshold of 0.70, and HMDB failed to annotate any correct compound with the same score threshold. We also performed annotations for standards with MetFrag CLI version 2.5.0 and PubChem [60] as the search database. Table 4 gives an overview of the annotations obtained by MAW, Spectra, SIRIUS, and MetFrag.

**Table 4:** Overview on the annotated features in MAW, SIRIUS, Spectra, and MetFrag, using the standards dataset, which had in total 37 features (and 9 standards). Total structural annotations refer to the percentage of annotated compounds that are the same in MAW, Spectra, and SIRIUS. Thus, 75.7% of the annotation in MAW were observed in Spectra and 70.2% of the annotations were observed in SIRIUS while some of the features had annotation sources from both Spectra and SIRIUS. We additionally ran MetFrag and among the top 100 candidates from each feature, 12 features had the same correct annotation. For the chemical classification, MAW annotated 97.3% of the features with a chemical class and all features had a molecular formula prediction. SIRIUS identified the chemical class (with CANOPUS) and molecular formula for all features, while Spectra and MetFrag did not provide these functionalities.

| Annotation level | MAW | Spectra (GNPS, HMDB, MassBank) | SIRIUS | MetFrag (PubChem) |
| --- | --- | --- | --- | --- |
| Total structural annotations | 100% (37) | 75.7% (28) (within MAW) | 70.2% (26) (within MAW) | 32.4%(12) |
| Chemical class prediction | 97.3% (36) | - | 100% (37) | - |
| Formula prediction | 100% (37) | - | 100% (37) | - |

### 3.3 Case study with the bryophytes dataset

#### 3.3.1 Marchantin compounds with the COCONUT database

MAW was used to annotate chemical structures from the spectra obtained from nine different species of the bryophyte dataset. As the species *Marchantia polymorpha* (group Marchantiophyta) is known to produce very specific cyclic bis-bibenzyls such as Marchantins and semi-cyclic Perrottetins which provide a challenge for computational annotation tools due to their cyclic structures, these compounds were chosen to evaluate the performance of MAW. In the ChemONT ontology, these compounds are represented by the compound superclasses Lignans, neolignans and related compounds or Stilbenes [61]. In the original study, three compounds were annotated as Marchantin compounds (Marchantin G, K, D). MAW annotated 4 Marchantin compounds and 1 Perrottetin compound within the top 25 candidates. Table 5 gives an overview of the annotated cyclic bis-bibenzyls in the *Marchantia polymorpha* samples.

**Table 5:**
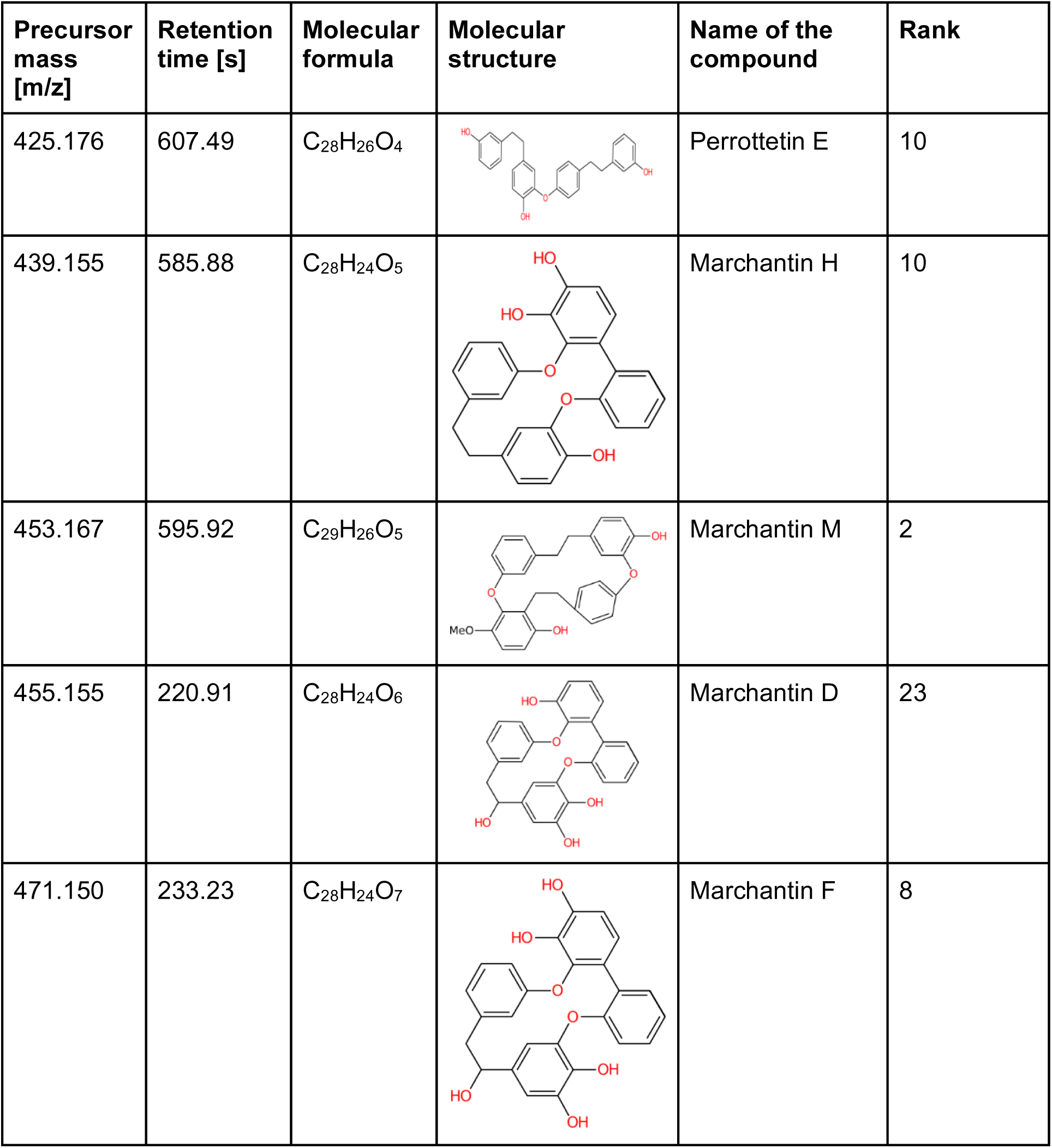
List of annotated cyclic bis-bibenzyls using MAW. The precursor mass [m/z] and retention time [s] represent the mass of the precursor ions given in mass-to-charge ratio and the median retention time of the precursor ion given in seconds, respectively. The molecular formula and name of the compound are extracted using SIRIUS. The chemical classes of the compounds are extracted from CANOPUS. All the molecular structures are generated via CDK-Depict. The given ranks are from MAW, using the database COCONUT in the compound dereplication module.

| Precursor mass [m/z] | Retention time [s] | Molecular formula | Molecular structure | Name of the compound | Rank |
| --- | --- | --- | --- | --- | --- |
| 425.176 | 607.49 | C<sub>28</sub>H<sub>26</sub>O<sub>4</sub> |  | Perrottetin E | 10 |
| 439.155 | 585.88 | C<sub>28</sub>H<sub>24</sub>O<sub>5</sub> |  | Marchantin H | 10 |
| 453.167 | 595.92 | C<sub>29</sub>H<sub>26</sub>O<sub>5</sub> |  | Marchantin M | 2 |
| 455.155 | 220.91 | C<sub>28</sub>H<sub>24</sub>O<sub>6</sub> |  | Marchantin D | 23 |
| 471.150 | 233.23 | C<sub>28</sub>H<sub>24</sub>O<sub>7</sub> |  | Marchantin F | 8 |

#### 3.3.2 Flavonoid “Nicotiflorin” as a highly probable annotation with spectral databases

Here, we describe the annotation and evaluation of the flavonoid Nicotiflorin using MAW. This compound was annotated by MAW in all of the bryophyte samples. The feature ID is file_1M287R318ID319 where File_1 is the file ID, M287 represents the precursor mass [m/z] and R318 represents the median retention time [s]. ID is an individual number assigned to the features in File_1. For ease of understanding, we will refer to this feature as M287R318. The detailed features are provided in Table 6.

**Table 6:**
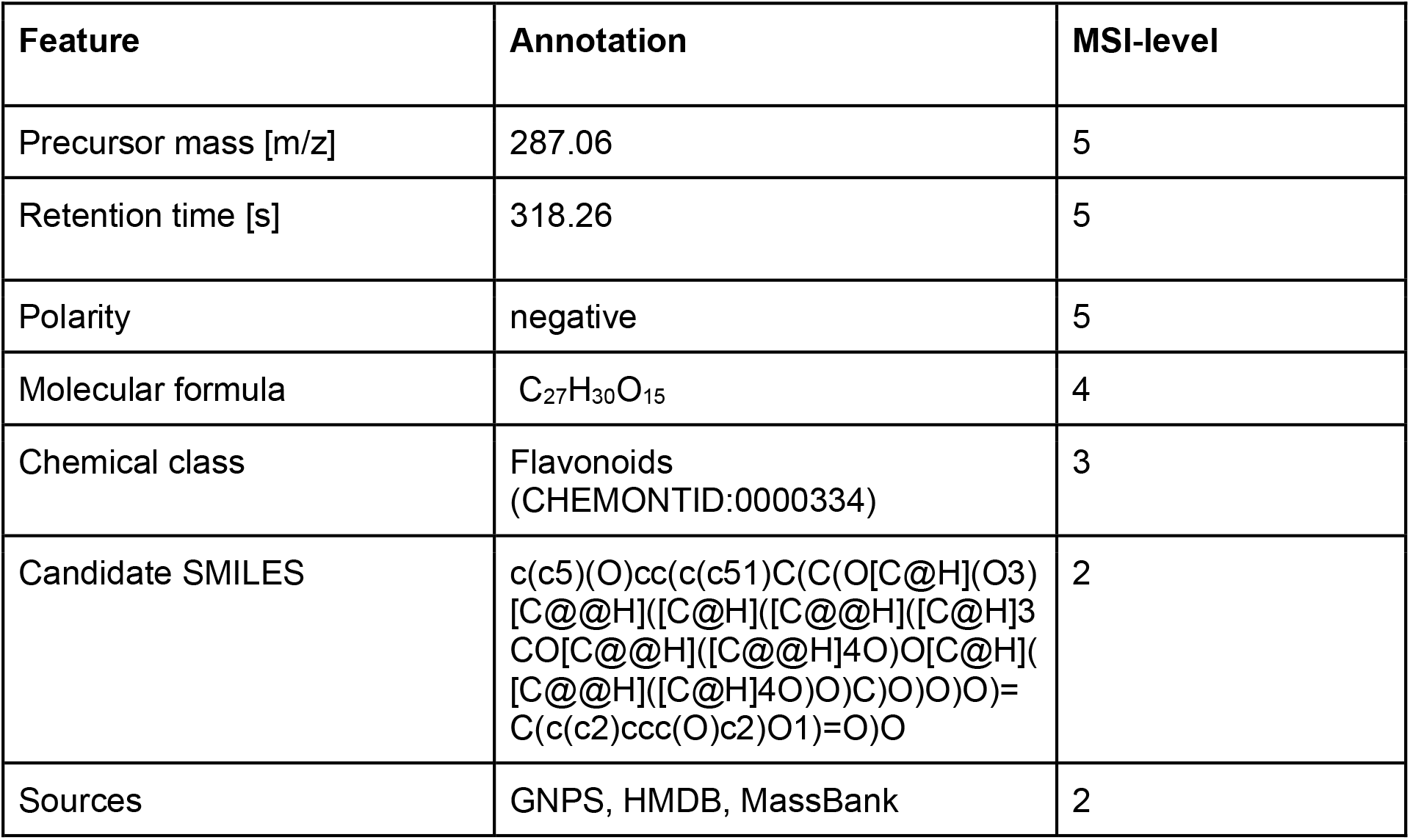
Features and annotations for feature ID M287R318 with MSI-levels. The measured precursor ion for Nicotiflorin has a mass [m/z] of 287.056. The retention time is given in seconds. This precursor ion was measured in negative mode during LC-MS. The molecular formula is extracted from PubChemPy and the class is annotated using ClassyFire. SMILES represent the structure of Nicotiflorin which was the same structure annotated with GNPS, HMDB, and MassBank.

MAW provides annotation results in two ways: (1) to generate a chemical similarity network between all top-scoring candidates, and (2) to select only one highly probable candidate for each feature.

For (1), MAW generates a TSV file, for each feature, that contains the nodes as candidates and Tanimoto similarity scores as edges. Figure 5 shows the chemical similarity network of candidates for M287R318 created with Cytoscape, importing the TSV file as a network in Cytoscape. The TSV file defines the start and end nodes (candidates) and their structural similarity represented by the Tanimoto similarity score. This visual representation helps in candidate selection by choosing the cluster with the highest number of annotation sources (GNPS, MassBank, HMDB, SIRIUS) and the highest-ranking candidate from these sources.

**Figure 5:**
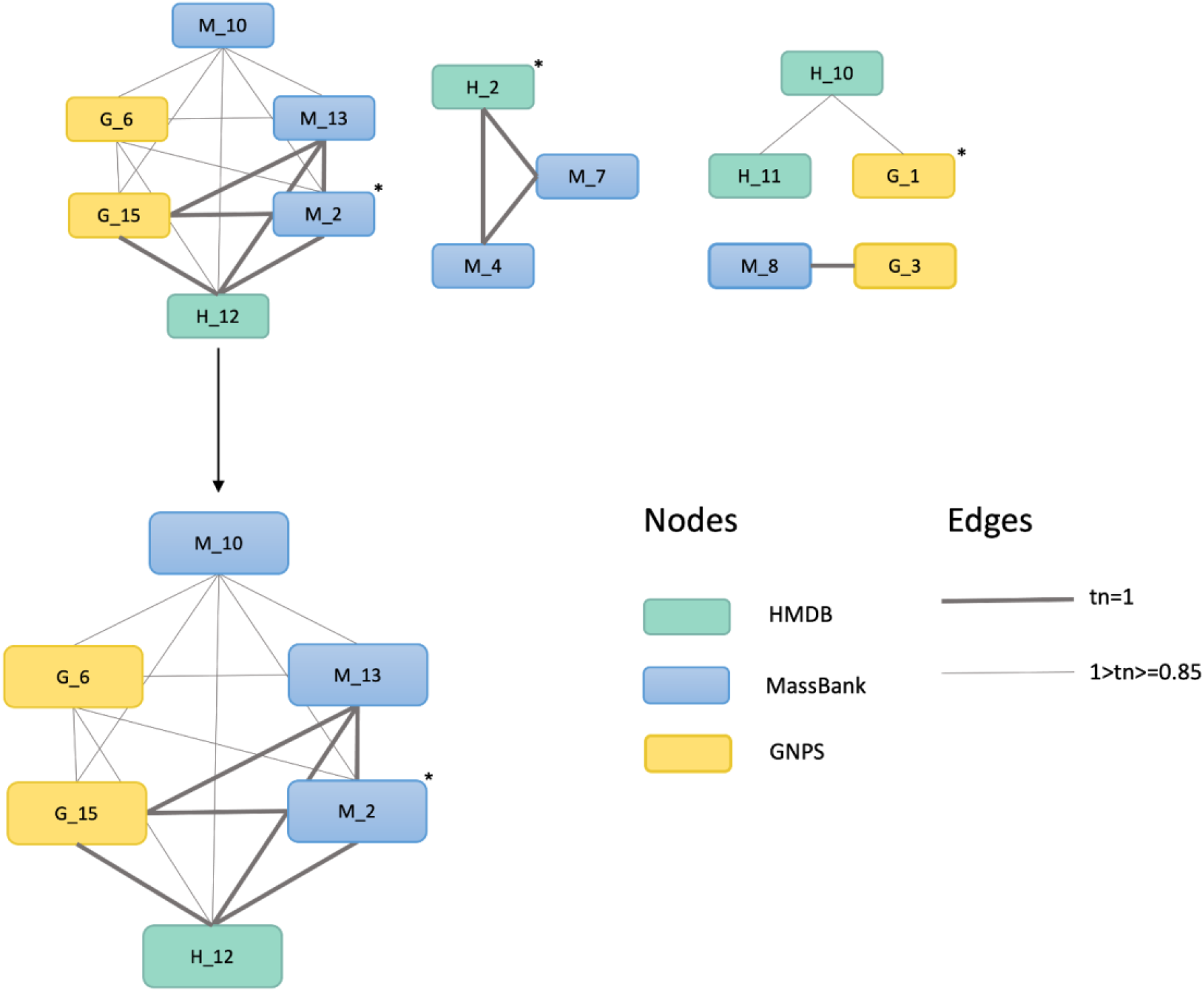
Chemical similarity network depicting the structural similarities among the top candidates from GNPS, MassBank, and HMDB (no annotations from SIRIUS are recorded for this feature). The nodes represent a chemical structure (as candidates). The edges represent the Tanimoto similarity score (tn) between the two nodes. The asterisk (*) shows the top-ranking candidate from each source that is present in the network. The network with the most nodes is zoomed in to visualize that G_15 (15th rank candidate from GNPS), H_12 (12th rank candidate from HMDB), M_2, and M_13 (2nd and 13th rank candidates from MassBank) have a structure with a Tanimoto similarity score equal to 1 and hence it is most probable to consider this structure to be annotated to the feature M287R318.

For (2), the chemical similarity network selects the most probable cluster which in this case is presented in Figure 5. The cluster represents three sources (spectral databases). This gives a higher probability to the structure in that cluster and can be assigned as the final candidate, in this case, Nicotiflorin. Using the package Spectra, a mirror image of MS^2^ spectra, both from the query and the top candidate from either of the spectral databases, can be created.

## 4 Discussion

Our Metabolite Annotation Workflow (MAW) facilitates and automates the complex and often cumbersome annotation of untargeted tandem mass spectrometry data in large metabolomics studies. To this end, MAW also improves candidate selection by integrating spectral and compound libraries and databases with computational annotation tools such as SIRIUS. It is also among the first tools that provide whole metabolome annotation and assess the chemical diversity of the biological samples.

### 4.1 Evaluation of MAW

Many software suites and R packages have been developed in the last years to solve the metabolomics annotation challenge, but it still remains the “bottleneck” in metabolomics due to an “infinite chemical dark space” [62]. The Metabolome Annotation Workflow (MAW) is specifically developed to aid the annotation process by automation of candidate selection within this space.

Computational tools such as MetFrag, SIRIUS(CSI:FingerID), and CFM-ID (Competitive Fragmentation Modeling for Metabolite Identification), generate *in silico* spectra of chemical structures from different compound databases based on their specified algorithms. This *in silico* spectral matching provides an annotation of up to MSI-level 3 [12], which refers to a tentative structure, not yet confirmed with a spectral library match or a standard (see Table 1). To provide more confidence in the annotation, the input spectra should be screened against actual experimental spectra. For this purpose, some tools and R packages have been developed such as Spectra, matchms [63], MS2Query (specifically for Analogues), or GNPS MASST, which use modified cosine similarity score or deep learning-based scores to match the MS^2^ query spectra against spectral databases. However, in general, there is a huge mismatch between submitted experimental spectra to spectral databases and chemical structures present in compound databases. This leads to a low number of annotated features. As MAW combines both *in silico* generated spectral matching and experimental spectral matching to increase the number of confident annotated metabolites, the percentage of annotated metabolites with MAW is higher than with using spectral or compound databases alone (see Table 4).

Another advantage of MAW is the automated candidate selection which allows one to choose the most optimal results. Generally, annotation tools provide a list of ranked molecules based on the associated scoring schemes and the user has to manually go through these candidates or just take the first candidate. This requires a lot of time and effort with big datasets. MAW not only provides these ranked lists but also provides a concise list of highly probable candidates from different databases automatically. The structural similarity between these candidates can be visualized in a chemical similarity network. A function to get only one candidate per feature is also provided. All the intermediate results are kept in an efficiently designed directory system, so the user can always refer back to the origin of the candidate. This way provenance records are stored for a given dataset, as the workflow is executed.

A major component behind the candidate selection is the scoring scheme followed throughout the workflow. We used the scoring schemes provided by Spectra and SIRIUS (CSI:FingerID), and the Tanimoto similarity score for the structural matching. Spectra uses the most common spectral matching metric which is the cosine similarity score that calculates how much overlap is observed between two spectra [64]. For HMDB and MassBank, this metric is used unaltered. Some good candidates are missed due to lower scores with high spectral matching as in the case of butanoyl carnitine and MassBank. For GNPS, this similarity score is altered as described in the method section and the spectral matching with GNPS gives better results as compared to the other two spectral databases. We conclude that spectral matching needs better metrics. The possible cases where cosine similarity metrics fail could be due to different fragmentation patterns under different experimental setups, or if there is a shift in the fragmentation peaks, the overlap between matching spectra would also be shifted [65]. To deal with this problem in a better way, we also consider the individual fragment mass [m/z] and intensity overlap, also counting in the number of matching peaks. All of these MAW-defined metrics should at least have a 0.5 matching score in addition to a high cosine similarity score. In this way, we could lower the possibility of getting any false positives but could end up missing candidates with low cosine similarity scores as false negative candidates. This could be solved by integrating Spec2Vec into MAW. Spec2Vec is a deep learning-based spectral matching score and has shown better spectral matching results as compared to simple or modified cosine similarity scores [65]. In addition, the public spectral databases are not completely standardized and hence can only annotate spectra present in the databases such as in the case of betaine, where reduced state trimethylglycine is annotated.

To further reduce the false negative results from spectral database screening is to use compound databases to compensate for all the known compounds which were not assigned a candidate from spectral databases. Here, MetFrag would be a good option when using a more concise library of molecules. Larger databases such as PubChem can lead to a larger list of candidates with the actual structure being at a lower rank (e.g: top 100th candidate), such as seen with candidates from the standards dataset, explained in Table 4. For the first version of MAW, we have only integrated SIRIUS (CSI:FingerID) for compound databases, a state-of-the-art annotation tool [66] into MAW. For features with no structural annotations, SIRIUS and CANOPUS provide a molecular formula and a chemical class. Thresholds for ranking SIRIUS candidates were considered, but SIRIUS CLI doesn’t provide a deterministic probability score, as a result, providing a threshold would differ from case to case, and hence the idea was discarded. In addition, the annotation of lipid classes using SIRIUS can lead to false positives due to the complex nature of lipids, and many lipid isomers co-elute in mass spectrometry, making it a challenge to annotate lipids using *in silico* libraries [67].

The annotation result evaluation for MAW has been accomplished via the standards dataset and the LC-MS^2^ data from the bryophytes datasets. MAW performed better for the standards dataset, keeping the actual compounds within the top three candidates. These standards are usually acquired with the LC-MS^2^ technique and have been submitted to the spectral databases. As a result, the experimental spectral matching led to the true positive as the top first annotation in most cases. For the annotation results from the bryophytes dataset, we could only annotate 8 compounds that were the same as in the original study. All the annotations from the original study were performed manually and most of the annotations were not within the top-ranked candidates. The main objective of the study was to identify the chemical classes and the annotation of natural products from different bryophytes species. Despite the fact that bryophytic organisms received much attention regarding their unique diversity of natural products, many features remain without an annotated chemical structure.

Additionally, some classes of natural products from these organisms are experimentally challenging to acquire with tandem mass spectrometry [68]. Using databases specifically focusing on natural products like COCONUT and GNPS can increase the probability of finding the compound of interest within the top-ranked candidates. We observed that using the “bio” database from SIRIUS, the actual compounds were ranked lower as compared to using the COCONUT database. Hence, it is important to have an objective for any annotation experiment, whether it is to perform targeted LC-MS to confirm the presence of known compounds, or it is to annotate novel natural products. The usage of different databases can lead to different results for less-studied species and compounds. As seen during the annotation for the chemical compounds from the Marchantin group, these compounds were ranked low due to their macrocyclic structures; the spectra for such structures are already challenging to obtain due to complex fragmentation patterns [69]. In such cases, it helps to have a pre-defined suspect list even in the case of an untargeted study.

The implemented functionalities enable MAW to provide an overview of the chemical diversity present in any dataset. MAW also can generate graphs to visualize the chemical similarities in a network or the chemical classification. All the steps in MAW are customized specifically for the annotation step of the metabolomics workflow. MAW also complies with the FAIR guidelines, making it Findable and Accessible on GitHub as an open-source code repository, and access to all the datasets, and libraries used for the development of MAW is provided via Zenodo. MAW is made interoperable as it is executed inside docker containers on any system. It also follows the Reusability principle by tracking all the intermediate results and collecting the provenance of the workflow execution.

### 4.2 Future Developments

To enhance spectral database matching, we recommend MAW to be integrated with Spec2Vec and matchms, along with MS2Query for providing structurally similar analogs. To avoid wrong predictions with candidates that don’t have a natural product like structure, a functionality that removes any rare natural product elements can be added. To enhance the automated candidate selection, we are also aiming to add MetFrag with PubChemLite and COCONUT. We are also planning for the support of a wider database selection in SIRIUS 5. The spectral similarity-based classical molecular network (MN) is another important aspect for the annotation of metabolites, among single-condition datasets or between two conditions, and linking these MNs to pathway annotations. To further support the FAIR principles, our idea is to extend the specification of the workflow in the Common Workflow Language (CWL) [70], gathering detailed provenance of workflow executions, and adding rich metadata about the workflow, its components, and the data used and produced by it.

## Conclusion

Structural annotation remains a complicated and time-consuming process. MAW is an automated metabolite annotation workflow. Using both spectral and compound databases, MAW helps to annotate the query spectral data to a corresponding experimental tandem mass spectra or to an *in silico* generated corresponding spectra in case there are no associated spectra present in spectral databases. Results of annotated metabolites from MAW can be further functionally mapped to metabolic pathways to aid in the structure prediction of compounds and can be integrated with other omics data to understand the biological and ecological significance of these metabolites. This could support the chemical characterization in diverse fields such as biomarker discovery in clinical metabolomics, and defining the chemical diversity of ecosystems, leading to the identification of novel natural products.

## List of abbreviations

CFM-ID: Competitive Fragmentation Modeling for Metabolite Identification
CLI: Command Line interface
COCONUT: COlleCtion of Open Natural ProdUcTs
CWL: Common Workflow Language
DDA: Data Dependent Acquisition
FAIR: Findable, Accessible, Interoperable, Reusable
GNPS: Global Natural Products Social Molecular Networking
HMDB: Human Metabolome Database
LC-MS: Liquid Chromatography Mass Spectrometry
LC-MS^2^: Liquid Chromatography-Tandem Mass Spectrometry
MAW: Metabolome Annotation Workflow
MAW-Py: Metabolome Annotation Workflow-Python Segment
MAW-R: Metabolome Annotation Workflow-R Segment
MCSS: Maximum Common Substructure
MSI: Metabolomics Standards Initiative
SMILES: Simplified Molecular Input Line Entry Specification
W4M: Workflow4Metabolomics

## Declarations

### Availability of data and materials

**MAW:**

- MAW can be installed with all its functions and dependencies via two docker images submitted to DockerHub [37, 38].
- Detailed Tutorial for MAW is available on GitHub Wiki, with associated Jupyter Notebooks present on GitHub [36].
- The Source Code is available on GitHub and MAW 1.0.0 version is available on Zenodo with the DOI: 10.5281/zenodo.7148450. An updated version (MAW 1.0.1) is present with the DOI: 10.5281/zenodo.7215518
- All the scripts used to run MAW on diatom and bryophyte datasets and scripts used to download databases are available on MAW-Bechmark repository (https://github.com/zmahnoor14/MAW-Benchmark) [71].

**MS Datasets**

- Standards: The LC-MS^2^ standards data generated by Vera Nikitashina are present on the MetaboLights repository under accession MTBLS3177(still under curation). These standards are also uploaded to Zenodo as mzML files with DOI: 10.5281/zenodo.7106205, which were used in MAW.
- Bryophytes: The bryophytes dataset is also available on MetaboLights under accession MTBLS709. The MS^2^ data file used in this study is available on Zenodo with DOI: 10.5281/zenodo.7107096
- Spectral Databases as Rdata: The currently available versions of the GNPS, HMDB, and MassBank are present on Zenodo with DOI: 10.5281/zenodo.6528931

**Programming language:** R 4.2.0 and Python 3.10.7

**Requirements:** Docker

## Authors’ contributions

MZ developed Metabolome Annotation Workflow (MAW), wrote the manuscript, made MAW and relevant data findable, accessible, and reproducible, and modified Docker images.

LG conceptualized and made MAW FAIR, made Docker images, parallelized R functions, and revised the draft.

CS supervised the project, obtained the funds, and revised the draft. MS conceptualized the project, supervised, and revised the draft.

KP provided the bryophytes dataset, supervised the project, and revised the draft. All authors reviewed the manuscript.

## Acknowledgments

The authors would like to acknowledge:

**Georg Pohnert (ORCID: 0000-0003-2351-6336) Lab** (Friedrich Schiller University Jena) for providing the data for the standards dataset, especially **Vera Nikitashina** (Friedrich Schiller University Jena)

**Ivo Große Lab** (Martin-Luther-Universität Halle-Wittenberg) for initiating the idea for candidate selection

**Kohulan Rajan (ORCID: 0000-0003-1066-7792)** (Friedrich Schiller University Jena) for the MAW logo, cleaning code, and helping with setting up the GitHub repository.

**Adelene Lai (ORCID: 0000-0002-2985-6473)** (University of Luxembourg, and Friedrich Schiller University Jena) and **Jonas Schaub (ORCID: 0000-0003-1554-6666)** (Friedrich Schiller University Jena) for ideas on candidate selection function

